# Spontaneous and automatic processing of magnitude and parity information of Arabic digits: A frequency-tagging EEG study

**DOI:** 10.1101/2019.12.26.888651

**Authors:** Mathieu Guillaume, Alexandre Poncin, Christine Schiltz, Amandine Van Rinsveld

## Abstract

Arabic digits (1-9) are everywhere in our daily lives. These symbols convey various semantic information, and numerate adults can easily extract from them several numerical features such as magnitude and parity. Nonetheless, since most studies used active processing tasks to assess these properties, it remains unclear whether and to what degree the access to magnitude and especially to parity is automatic. Here we investigated with EEG whether spontaneous processing of magnitude or parity can be recorded in a frequency-tagging approach, in which participants are passively stimulated by fast visual sequences of Arabic digits. We assessed automatic magnitude processing by presenting a stream of frequent *small* digit numbers mixed with deviant *large* digits (and the reverse) with a sinusoidal contrast modulation at the frequency of 10 Hz. We used the same paradigm to investigate numerical parity processing, contrasting *odd* digits to *even* digits. We found significant brain responses at the frequency of the fluctuating change and its harmonics, recorded on electrodes encompassing right occipitoparietal regions, in both conditions. Our findings indicate that both magnitude and parity are spontaneously and unintentionally extracted from Arabic digits, which supports that they are salient semantic features deeply associated to digit symbols in long-term memory.

---

Number symbols convey a large range of different semantic properties, such as being a prime number, being a multiple of another number, or belonging to the power of another number. While these various properties need to be assimilated by long and laborious cultural learning processes, some of them are critical for understanding number symbols’ essence ^[1]^. Among these properties, cardinality (or numerosity) and parity are of crucial interest because they are both acquired at an early stage of numeracy learning and they are known to increasingly become salient properties of number symbols ^[2:7]^. The question whether and to what extent such semantic number properties are automatically activated has occupied the numerical cognition literature since several decades ^[8:22]^. This issue is essential because it directly questions the nature of semantic representation of numbers, as well as the developmental trajectory of numerical and mathematical learning ^[23:29]^. For instance, 3^rd^ graders typically associate digits based on magnitude characteristics when they are asked to categorize numbers according to similarity, while 6^th^ graders are more influenced by parity; adults, for their part, equally categorize digits on their parity and magnitude features ^[30]^.

## Automaticity in magnitude processing

Automaticity is usually defined as an unconscious and unintentional process, that might occur in parallel with other processes ^[31]^, and that happens without monitoring ^[32, 33]^. From a general cognitive perspective, automatic (and intuitive) processes are generally contrasted to slow logical reasoning processes ^[95:98]^. Despite this duality, it has been argued that a clearer distinction should be made within automatic processes, between two different types of automatic processes ^[21, 32, 34]^. The first automatic process type is intentional, used when it is necessary and beneficial for the ongoing task, for instance manipulating a number during calculation or reading words in a sentence. The second type is autonomous, unintentional and used even when its processing is irrelevant to the ongoing task. It is noteworthy that pointing out unintentional automatic processing from intentional processes is experimentally complicated. It is actually not possible to disentangle a useful and intentional process from an autonomous process when this process is required for the task ^[21]^.

Despite of these difficulties, three effects could depict automatic processing with number magnitude. The first is the *SNARC effect* (Spatial Numerical Association of Response Codes) which is characterized by an influence of the magnitude while participants are asked to categorize numbers according to their parity ^[34]^. In particular, participants are faster to categorize small numbers with their left hand and large numbers with their right hand independently of the number’s parity. Many authors explain this effect by suggesting the automatic activation of a mental number line ^[36]^ during the task, where the smaller numbers are mapped on the left and the larger numbers on the right^1^. Nonetheless, because parity is also a numerical dimension, experimental settings could trigger the automatic processing of magnitude. The triggering could happen when the relevant dimension (i.e., parity in SNARC effect) and the irrelevant dimension (i.e., magnitude in SNARC effect) are related. In other words, parity and magnitude are both numeric, thus it is difficult to know if magnitude activation observed during the SNARC effect is an autonomous or an intentional automatic processing^2 [21]^.

A second domain revealing automatic number magnitude processing refers to the *interference effects* observed in tasks similar to the well-known Stroop experiment ^[51]^. One type of such numerical interference effects is for instance the semantic congruity effect. Adults are indeed faster to judge the physical size of two numbers when the bigger number was also the larger in terms of numerical magnitude (e.g., comparing **5** vs. 2 is easier than 5 vs. **2**) ^[13, 22, 25, 52, 53]^. Another example is the size Congruency Effect happening when participants are asked to choose between two displays with digits, the display with the more numerous array of digits. Participants are faster during this task when the display comprising more digits also contains numerically larger digits and the screen with fewer digits also contains numerically smaller digits ^[18]^. However, these interference effects are only observed in numerical tasks and consequently numerical magnitude could be triggered by the task itself ^[21]^. According to this line of reasoning, interference effects are thus not sufficient to argue that number magnitude is accessed automatically in an autonomous manner.

Third, there is the distance effect, which refers to the fact that comparing close numbers (e.g., 4 and 6) is harder than comparing more distant numbers (2 and 8). In other words, the difficulty of comparing two numbers is inversely related to the numerical distance between these two numbers ^[54]^. Distance effects are found when participants are explicitly instructed to compare numbers ^[11]^, but also when participants had simply to say if two Arabic numbers were the same or not ^[13]^. Nonetheless, such evidence has also been assumed to be insufficient to accept the distance effect as a proof of autonomous automatic processing ^[21]^. That is to say, even if the distance effect occurs in non-numerical tasks, processing the magnitude in this kind of tasks would be beneficial for the task performance. Distance effects could therefore be a marker of intentional automatic processing rather than marker of autonomous automatic processing.

## Automaticity in parity processing

Regarding parity, a number can only belong to one out of two categories: ‘*even*’ or ‘*odd*’, indicating that a number is divisible by two or not. Parity judgments are typically learned relatively early, around the two first years of formal education in western cultures ^[50]^, and even earlier in eastern cultures ^[55]^. While some authors argued that parity is computed by dividing a number by two ^[56]^, others proposed that parity information is stored as declarative memory content ^[35, 57]^. According to the latter, parity access could not be the result of computations because there is no size effect in parity tasks. More precisely, if the determination of parity were up to dividing by two, small numbers should be processed faster than large numbers, which is not the case. In the same vein, certain digits with specific properties are categorized faster as odd or even ^[4]^. This is for instance the case for numbers that are a power of two (i.e., 2, 4 and 8) and for prime numbers (i.e., 3, 5 and 7). In particular, the more specificities a given number has, the faster their parity will be accessed ^[35]^. These findings are in line with a recent study proposing that the objective, mathematically defined parity differs to a certain degree from the “perceived” parity, as some numbers seem to be more prototypical for their categories ^[58]^.

To the best of our knowledge, there are only a limited number of investigations concerning the automaticity of parity processing. A study of parity categorization using subliminal priming of Arabic numbers (i.e., participants could not consciously see the primed digits) before the onset of a given target number word showed that participants were faster to categorize the target when the primed digit and the target number had the same parity ^[59]^. These results are in favour of an automatic access to the parity information of digits. This is in line with other studies that assessed arithmetical fact verifications based on the parity of the multipliers or addends ^[60]^. In particular, those tasks consisted in judging the correctness of additions or multiplications facts. If the parity rules (such as: addition or multiplication of two even numbers gives an even number) were not respected, participants were faster to say that arithmetical facts were false ^[57]^. Some authors argued that the application of the parity rules is unconscious and automatic because some of their participants reported that they were unaware of using the parity rules, while their results indicated its utilisation ^[60]^. Nevertheless, in both previous paradigms, the experimental task required direct manipulation of the number symbol and therefore can only point towards intentional automatic processing ^[21]^, leaving open the question whether autonomous unintentional automatic parity processing might also arise in some circumstances.

## Neural correlates of (automatic) number processing

Neuroimaging studies consistently indicated that the parietal cortex and more specifically the intra-parietal sulcus (IPS) plays a pivotal role in the processing of numerical information ^[61:65]^. Indeed, this region has been associated to a various range of tasks involving the manipulation of numerical material ^[66, 94]^. Magnitude comparison of number symbols activate more predominantly the right IPS, for conjoint symbolic and non-symbolic comparison ^[92]^, but also for exclusively symbolic comparison ^[64, 91]^. Moreover, the same bilateral IPS region was activated during numerical magnitude comparison as well as parity categorization in a PET study ^[63]^, in which the authors consequently proposed that parity and magnitude are processed in neighbouring areas. It was also proposed that both the occipitoparietal network used for perceptual and representational processing of Arabic digits and the frontoparietal network used for semantic processing are involved in numerical processing such as magnitude comparisons and arithmetical facts retrieval ^[64, 65]^.

The parietal cortex is also associated with automatic processing of number symbol semantics. Tasks requiring the simple detection of visual numerals or number words activate the IPS significantly more than the same tasks with non-numerical stimuli (*e.g*., detection of letters or colours ^[69]^). To the best of our knowledge, only a few studies further investigated automatic responses to number symbol magnitudes, and none examined automatic parity processing. Noticeably, using fMRI, a study reported stronger BOLD responses for large digits (8 or 9) than for small digits (1 or 2) within two bilateral parietal regions (i.e., the inferior parietal lobule and the IPS), when investigating the neural correlates of non-predictive number cues ^[70]^. Other studies reported greater ERP amplitudes across parietal electrode regions for small digits compared to large digits, in a task where numbers were used as non-informative attentional cues ^[71, 72]^. Nonetheless, imaging evidence in favour of unintentional processing of magnitude or parity is currently lacking.

## Current study and experimental design

In the present study, we aim at assessing the autonomous unintentional automaticity of magnitude and parity processing, carefully avoiding any triggering of numerical information processing (such that the existence of autonomous unintentional automaticity can be probed ^[21]^). For that purpose, we used a sensitive frequency-tagging approach – Fast Periodic Visual Stimulation (FPVS) – to record electrophysiological responses tagged at the frequency of experimentally manipulated magnitude and parity changes, in a passive viewing setting. Similar FPVS paradigms relying on passive viewing have successfully been used to study high-level discrimination of faces ^[73]^, tools ^[74]^, as well as reading (word recognition ^[75]^, and letter strings discrimination ^[65, 76]^), and non-symbolic quantities^[77]^, without engaging participants in any kind of explicit processing of the presented stimuli.

We combined the FPVS paradigm with a standard/deviant procedure (*oddball* method ^[79]^). More precisely, during one-minute sequences, we displayed digits at the very fast rate of 10 Hz (ten stimuli per second) following a sinusoidal contrast modulation. Within the stream of standard digits, a periodic deviant was introduced every eight trials such that digits from the deviant category appeared at 1.25 Hz (see Figure 1). The digits used as standards or deviants varied according to the condition: In the *magnitude* condition, digits smaller than five were presented as standard and digits larger than five as deviants, or the other way around. In the *parity* condition, odd digits were presented as standards and even digits as deviants, or the other way around. Finally, in a *control* condition, half of the digits were arbitrary categorized as standards and the other half as deviants (see Figure 1).

**Figure 1.**
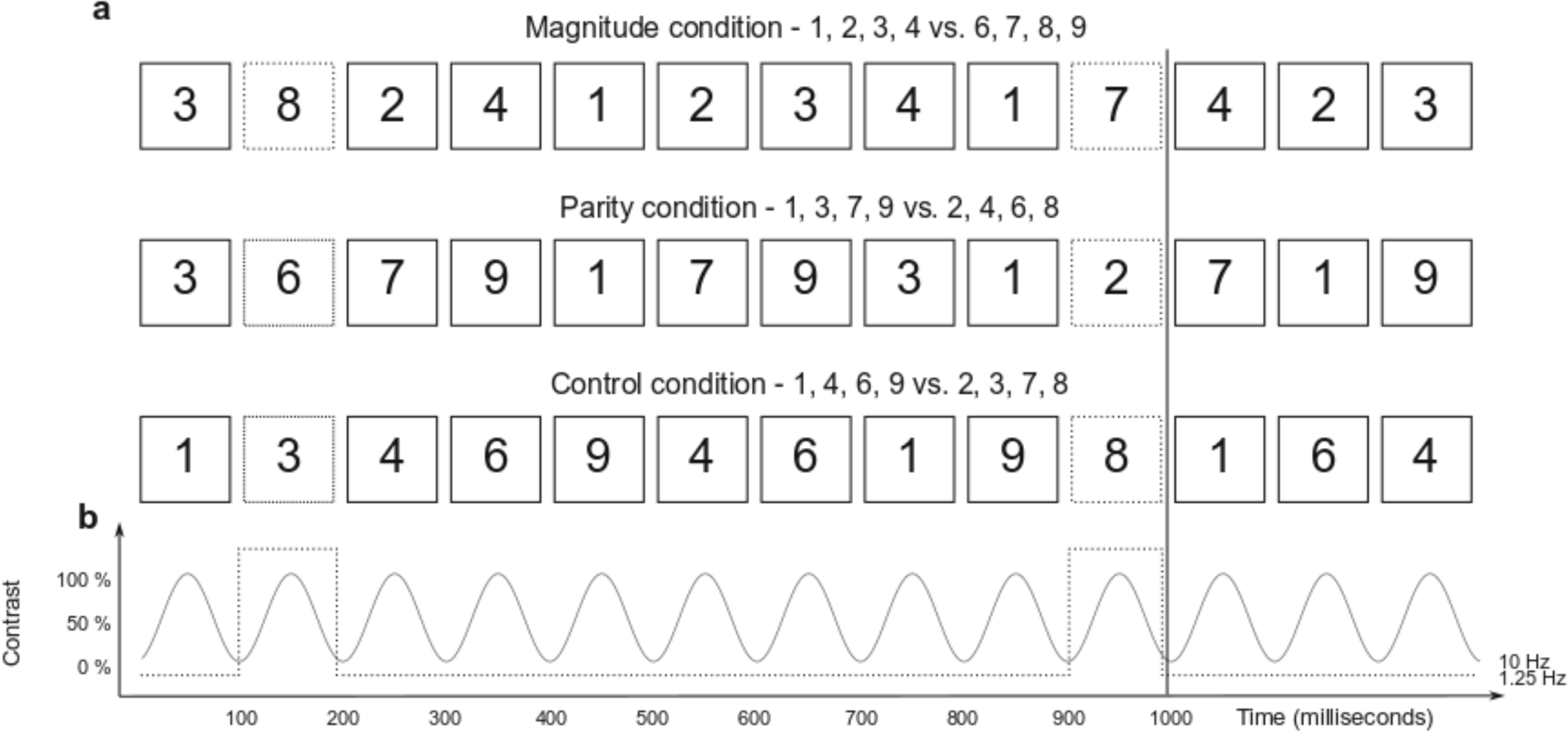
Illustration of the Fast Periodic Visual Stimulation paradigm. (a) During four 60 second sessions, Arabic digits between 1 and 9 (except 5) were periodically displayed at the base frequency of 10 Hz. In all conditions, digits from a determined standard category were periodically displayed at the base frequency, while digits from another deviant category were displayed at the oddball frequency of 1.25 Hz. In the Magnitude condition, categories were based on the magnitude of the digits (smaller or larger than five). In the Parity condition, categories were constructed as a function of the parity of the Arabic digit (odd or even). In the Control condition, categories were arbitrary. Fonts, font sizes, and positions are here constant for better readability. (b) Illustration of the onset and the offset of each stimulus following the sinusoidal contrast modulation from a 0 to 100% contrast.

We postulate that we should record an electrophysiological response synchronized at the deviation frequency (i.e., 1.25 Hz) if the brain is sensitive to the changes from one category to another. In other words, if magnitude and/or parity are processed in an autonomous unintentional automatic way, then the category changes should systematically elicit a specific brain response at the frequency of the changes for the respective condition(s), while no such response should arise during the control condition as the latter’s digit categories were arbitrary and not related to any semantic content. Doing so, we can objectively measure whether a given semantic number information is automatically and unintentionally processed by the brain although no numerical task was requested from the participant.

## Results

Figure 2a illustrates the topographical maps of the averaged oddball amplitudes for every condition, expressed as the Signal-to-Noise Ratio (SNR) of the frequency bin of interest and its harmonics up to the seventh (i.e., 1.25, 2.50, 3.75, 5.00, 6.25, 7.50, and 8.75 Hz). These maps highlight that the strongest recorded responses in each condition were located over a bilateral region encompassing occipitoparietal areas. It is worth noticing that there was a substantial overlap of the location of the brain responses between the experimental conditions, which allows direct comparison of amplitudes recorded within these regions.

**Figure 2.**
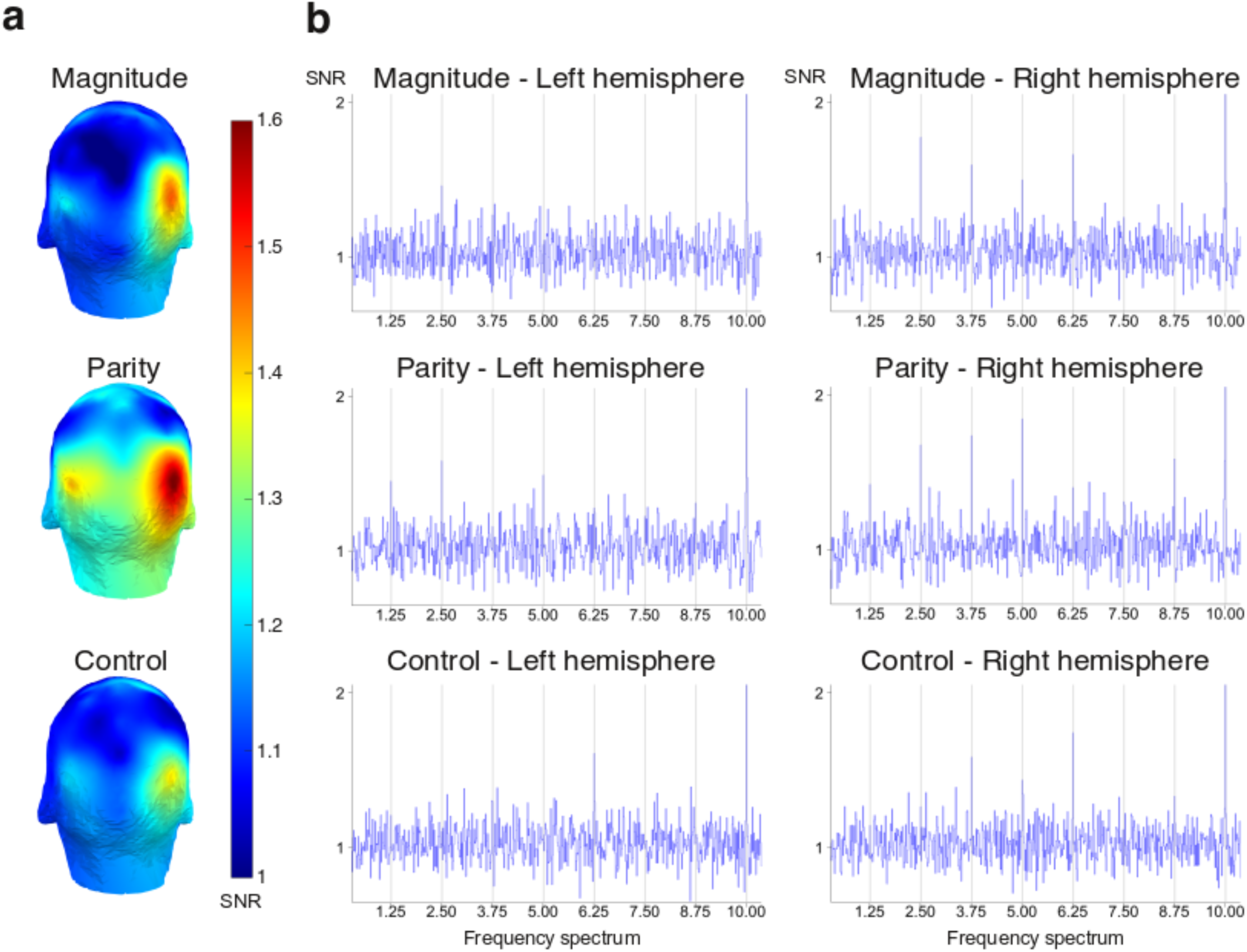
(a) Topographical maps of the periodic brain responses expressed as the Signal-to-Noise Ratio (SNR) of the average of the amplitude at the oddball frequency (1.25 Hz and its harmonics up to the 7^th^). (b) Amplitudes spectra expressed as SNR for the three conditions (magnitude, parity, and control) within the left electrodes of interest (A9, A10, A11) and within the right electrodes of interest (B6, B7, B8).

Figure 2b represents the EEG spectra recorded on the electrodes from the left (A9, A10, A11) and right (B6, B7, B8) occipitoparietal regions of interest corresponding to the topographical maps. The spectra, expressed as SNRs, are averaged across participants for every condition. In each condition, there were clear peaks at 10 Hz, with Z-scores systematically reaching values larger than five. These findings support that both occipitoparietal regions specifically synchronised to the 10 Hz base rate during the recording sessions. Note, however, that neural synchronisation to the base rate merely reflects brain response to the periodic stimulus onsets at 10 Hz.

More critically for the purpose of the current study, we also observed in Figure 2b large peaks in the frequency bins that are harmonics of the periodic category fluctuation (up to the seventh harmonic). In the Magnitude condition, there were fewer clear peaks across these frequency bins within the left electrodes, but peaks were clearly depicted in the right hemisphere. For the Parity condition, we observed some clear peaks within the left region of interest, and many definite peaks within the right region of interest. Finally, more unexpectedly, we also found some responding peaks in the control condition, mostly located within the right occipitoparietal regions.

In order to assess whether these peaks were statistically significant, we computed the average signal amplitude of the frequency of interest and its harmonics up to the seventh, expressed as a Z-score, as a function of the condition (see Figure 3). Within the three left electrodes of interest, mean oddball amplitude during the Magnitude condition was 0.51, *Standard Deviation (SD)* = 1.52, mean amplitude in the Parity condition was 1.59, *SD* = 1.26, and mean amplitude for the Control condition was 0.69, *SD* = 1.17. Within the right electrodes, mean amplitude during the Magnitude condition was 2.01, *SD* = 2.40, during the Parity condition it reached 2.50, *SD* = 1.85, and it was 1.23, *SD* = 1.90, for the Control condition. Visual inspection of Figure 3 reveals that the right occipitoparietal regions of interest were significantly sensitive to the periodic category changes during both experimental conditions, with z-scored amplitudes exceeding the unilateral 95% threshold of 1.64.

**Figure 3.**
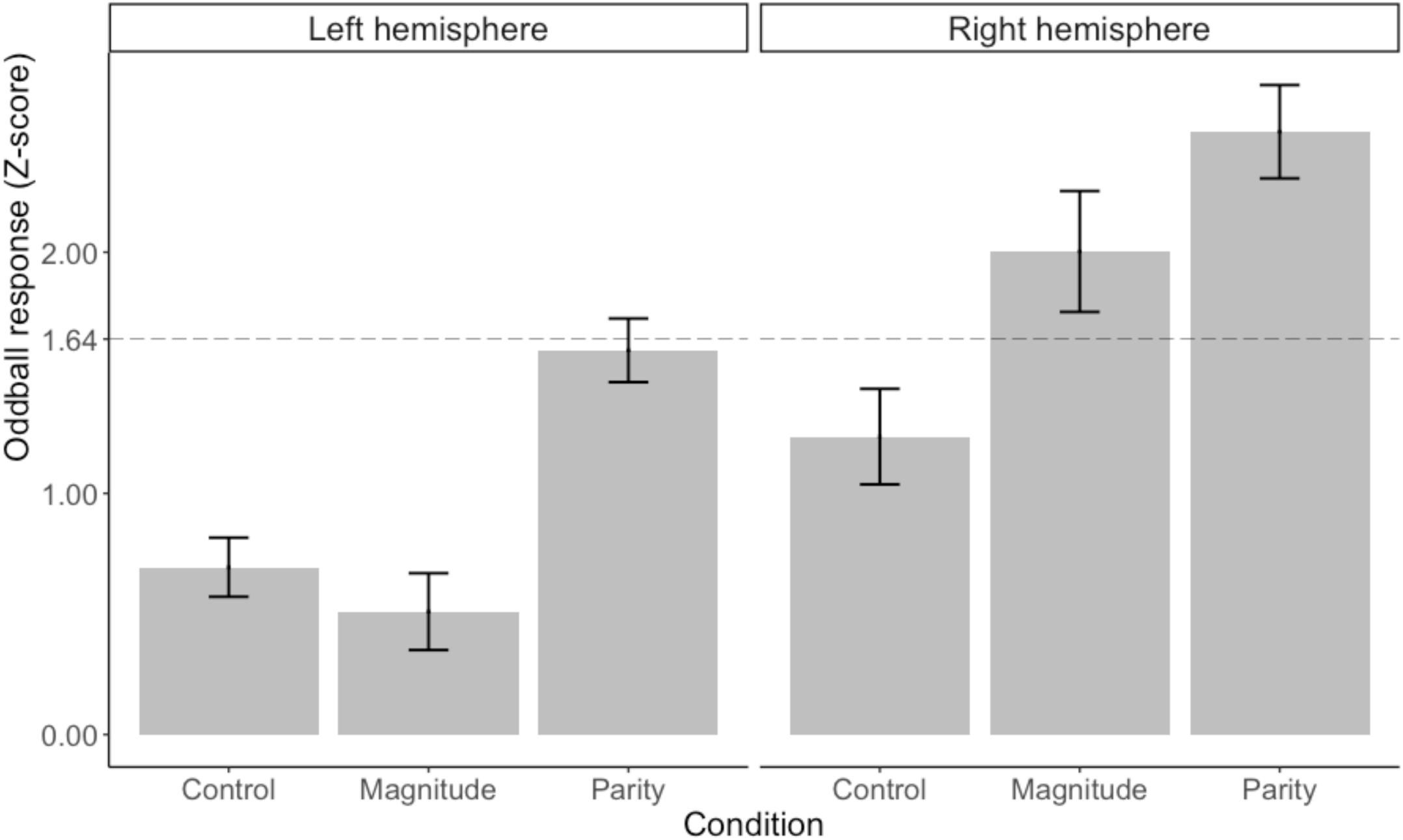
Mean amplitude, in Z-score, across the oddball frequency and its harmonics up to the seventh, as a function of the condition and as a function of the electrode lateralisation (left hemisphere: A9, A10, A11; right hemisphere: B6, B7, B8). Vertical bar depicts standard deviations. The horizontal dashed line represents the one-sided 95% threshold of significance (*Z* > 1.64, *p* < .05, one-tailed).

We additionally performed a repeated-measures ANOVA with the region of interest (two levels, either left or right) and the condition (three levels) as fixed factors. There was a significant main effect of the condition, *F* (2, 88) = 14.409, *p* < .001, partial η^2^ = .179, and a significant main effect of the region factor, *F* (1, 44) = 5.468, *p* = .024, partial η^2^ = .111. Although the interaction did not reach the significance level, *F* (2, 88) = 1.761, *p* = .178, partial η^2^ = .038, we conducted pairwise two-sided Wilcoxon tests (since assumption of normality was rejected) to directly compare the amplitudes within each hemisphere. These *post-hoc* tests revealed that, for the left electrodes of interest, Parity yielded significantly greater responses than both Control, *Z* = 56, *p* = .011, and Magnitude conditions, *Z* = 43, *p* = .003. On the other hand, Parity and Control did not differ from each other, *Z* = 168, *p* = .377. For the right electrodes of interest, Parity and Magnitude were significantly greater than the control condition, respectively *Z* = 35, *p* < .001, and *Z* = 65, *p* = .025, but the experimental conditions did not differ from each other, *Z* = 91, *p* = .160.

## Discussion

Magnitude and parity are very salient features of number symbols ^[5, 30]^. While there are several studies demonstrating the existence of automatic number symbols processing in educated adults, only a few studies could unequivocally demonstrate the existence of autonomous and unintentional number magnitude processing. Therefore, it is still necessary to provide further evidence by triggering magnitude as little as possible in order to disentangle autonomous from intentional automatic processing ^[21]^. Moreover, data concerning an automatic processing of parity are currently lacking.

The current FPVS paradigm allows investigating the spontaneous and autonomous nature of specific cognitive processes during passive viewing of rapid sequence ^[87]^. This technique has not yet been used to investigate number symbol processing, though it is very relevant for our objective since it is an objective measure of the brain response at a frequency defined a priori by the experimenter, which allows direct and straightforward analysis of the neural synchronisation. Moreover, the recorded responses quantify the processing of digits’ magnitude and parity without an active task and thus, without the implication of decision-making processes ^[78]^. Taken together, the current design avoided any conscious triggering of magnitude or parity during the recording session and thus probe the autonomous nature of these numerical processes.

We observed a significant neural synchronisation to changes in numerical magnitude and parity within right occipitoparietal areas, whereas no such synchronisation was found during the control condition. It is noteworthy that visual inspection of SNRs suggests that some brain responses peaked at some harmonics of interest in the control condition, where category was based on temporarily constructed rules, but these peaks did not reach the statistical significance. In contrast, the strong responses within the right occipitoparietal electrodes indicate that participants implicitly processed the category to which numerical stimuli belong and reacted to changes with respect to the category. Additionally, parity yielded significantly larger brain responses than the other conditions in the left hemisphere.

We thus measured substantial implicit responses to the two manipulated semantic features, but not to short-term arbitrary associations. This means that we can exclude that the brain synchronisations recorded in both experimental tasks were only due to the rare nature of the deviant stimulus. These evidences support the view that both numerical magnitude and parity were automatically accessed and processed during passive viewing sessions. In other words, both semantic features are spontaneously and unintentionally extracted from Arabic digits, which corroborates previous observations that several deliberate tasks on Arabic digits (such as naming) are necessarily semantically mediated ^[14]^. It is noteworthy that the automaticity of parity that we observed here does not support the hypothesis arguing that parity is computed by dividing the number by two ^[56]^, but our findings are rather in line with the idea of a direct retrieval from semantic memory ^[35, 57]^.

The topography of the frequency-tagged responses corresponds with seminal models of numerical cognition, such as the triple-code model ^[66, 80]^, since we found significant responses over right occipitoparietal electrodes specifically related to both magnitude and parity changes. This is in line with the hypothesis that the semantic of number representations are housed in parietal regions ^[98]^. The occipitoparietal responses that we obtained during Magnitude and Parity conditions are typical in ERP studies investigating the neural distance effect with Arabic numbers ^[64, 67, 68]^. These studies highlighted a neural distance effect illustrated by a modulation of the positivity of P2p recorded over occipitoparietal electrodes as a function of the numerical distance during numerical comparison. These studies, except ^[68]^, also found a dominance of the right hemisphere for numerical magnitude comparison. In the absence of an active task, we similarly found significant right-sided dominance for numerical symbols’ processing. We did not, however, observe any significant differences between right hemisphere electrodes responding to parity and magnitude, which is partly in line with a PET study indicating that parity and magnitude information are processed in neighbouring area inside the IPS ^[81]^.

Finally, it is noteworthy that each digit presentation only lasted 100ms in the current experimental design (since our base rate was 10Hz, see Figure 1). Previous ERP studies showed that number processing actually follows a time course that involves distinct stages of neural processes ^[99]^. Critically, numerical magnitude was already reported to be early extracted (within 200ms) from digit symbols ^[99]^. Due to the fast display period in the current FPVS paradigm, fewer than 100ms, we can assume that our method mostly captured early cognitive processes. Because we observed significant brain synchronization to magnitude and to parity, our results consequently support that not only magnitude, but also parity information, can be extracted and processed very early in the brain. The time course of the autonomous processing of numerical features could be investigated in future studies by, for instance, varying the base rate of similar FPVS paradigms. More generally, the current paradigm provides strong evidence of unintentional and autonomous processing of semantic information from digits, and it could be easily implemented to follow how symbols’ semantic processing becomes automatic across typical and atypical development.

In summary, we observed significant brain responses tagged at the frequency of magnitude and parity periodic changes over occipitoparietal electrodes during passive viewing of a rapid stream of Arabic digits. We demonstrate that numerical magnitude is spontaneously processed, and we also provide evidence that parity is a semantic feature that is unintentionally activated by Arabic digits in numerate adults. The current study thus provides evidence that culturally learned number symbols become automatically processed by the human mind.

## Method

### Participants

Twenty-nine undergraduate students from the University of Luxembourg participated in the study. Any history of neurological or neuropsychological disease or any uncorrected visual impairment constituted exclusion criteria. To ensure participants had no major mathematical difficulties, participants’ arithmetic fluency was evaluated with the Tempo-Test Rekenen ^[82]^, which is a timed pen-and-paper test (five minutes) consisting in arithmetic problems of increasing difficulty. All participants reached the inclusion criterion, which was 100 correct items out of 200, and were included into the present study. Six participants were excluded from the final sample due to substantial noise in their EEG signal (mostly due to transpiration). The final sample thus consisted in 23 adults, with a mean age of 24 years (*SD* = 3.5). Participants received 30 euros for their participation.

### Experimental setup

We used MATLAB (The MathWorks) with the Psychophysics Toolbox extensions ^[83:85]^ to display the stimuli and record behavioural data. The EEG recording took place in a shielded Faraday cage (288 cm × 229 cm × 222 cm). Participants were seated at 1 meter from the screen, with their eyes perpendicular to the centre of the screen (24’’ LED monitor, 100 Hz refresh rate, 1 ms response time). Screen resolution was 1024 × 768 px, with a light grey background colour. The order of the conditions during the EEG recording session was counterbalanced across participants.

### Material and Procedure

Although FPVS paradigm theoretically only involves passive viewing of the stimuli, we introduced a basic orthogonal active task during the recording sessions in order to ascertain that participants were looking at the computer screen. Participants were thus instructed to fixate the centre of the screen where a small blue diamond (12px size) was displayed, and they were asked to press a button with their right forefinger when they detected that the diamond changed its colour from blue to red. This colour change was not periodic and could randomly occur six to eight times in a given sequence. Participants were also informed that black digits ranging from 1 to 9 – excluding 5 – would quickly appear on the screen. They were explicitly instructed not to actively look at the digits but to keep their gaze on the central diamond. On average, participants took 640 ms (*SD* = 121 ms) to respond to the colour change that affected the fixation diamond. Misses were very rare, occurring only in 1.5% of the trials. Such high detection rate indicates that participants followed the instruction and kept their gaze on the centre of the screen during EEG acquisition.

Digits were sequentially presented at the fast base frequency of 10 Hz (*i.e.*, ten stimuli per second) following a sinusoidal contrast modulation from 0 to 100 % ^[73, 77, 86]^ (see Figure 1). There was thus for each stimulus a 50ms period of gradual fade-in and 50ms of gradual fade-out. The successive presentation of digits consisted in sequences that lasted 64 seconds, including 60 seconds of stimulation and 2 seconds of fade-in and fade-out, which we did not analyse.

We used the FPVS variation of the oddball design ^[79]^ in which we introduced a periodic fluctuation within the standard sequence: the stimulus category changed every eight items (*i.e.*, at 1.25 Hz). Our design included three different experimental conditions, each consisting in a different category change. In the *magnitude condition*, periodical variations were based on the magnitude of the Arabic digit (*i.e.*, smaller than 5 or larger than 5). For half of the sequences, the standard stimuli displayed at the base rate were randomly drawn among the smallest numbers (1, 2, 3, 4) whereas the deviant item was randomly drawn among the largest numbers (6, 7, 8, 9). For the other half, it was the opposite, the greatest numbers were standards and the smallest ones were deviant. In the *parity condition*, periodical variation was based on the parity of the Arabic digit (*i.e.*, odd vs. even). For half of the sequences, the odd category (1, 3, 7, 9) was standard and the even category (2, 4, 6, 8) was deviant, and vice-versa for the other half. Finally, we introduced a *control condition* that was based on two arbitrary categories with equal amounts of odd, even, small and large numbers (1, 4, 6, 9 vs. 2, 3, 7, 8). As the control categories were arbitrary, and in order to reduce the recording time, we only presented the first set as the standard category and the second set as the deviant category. Sequences for every condition were repeated four times. The magnitude and parity conditions each yielded eight sequences, for a total of twenty sequences.

Following previous recommendations ^[87]^, and to decrease habituation to the visual properties of the stimuli ^[9]^, we deliberatively introduced stochastic physical variations within the stimuli stream. The stimulus font randomly varied among four possibilities (Arial, Times New Roman, Cambria, and Calibri), the position of the stimulus randomly varied on both the vertical and horizontal axes (with a variation of maximum 10% from the centre of the screen), and the font size varied (from 122 to 148, with an average value of 135). These random visual variations occurred at each onset and were thus not congruent with our frequencies of interest (*i.e.*, 1.25 Hz and its harmonics).

### EEG acquisition

We used a 128-channel BioSemi ActiveTwo system (BioSemi B. V., Amsterdam, The Netherlands) tuned at 1024 Hz to acquire EEG data, as in ^[77]^. We positioned the electrodes on the cap according to the standard 10-20 system locations (for exact position coordinates, see http://www.biosemi.com). We used two supplementary electrodes, the Common Mode Sense (CMS) active electrode and the Driven Right Leg (DRL) passive electrode, as reference and ground electrodes, respectively. We held electrodes offsets (referenced to the CMS) below 40 mV. We also monitored eye movements with four flat-type electrodes; two were positioned lateral to the external canthi, the other two placed above and below participant’s right eye; but we did not further analyse these electrodes.

### EEG analysis

Analyses were conducted with the help of *Letswave 6* (http://nocions.webnode.com/letswave). Before starting the analyses, we down-sampled our data file resolution from 1024 Hz to 512 Hz for faster computer processing. We used a 4-order band-pass Butterworth filter (0.1 to 100 Hz) and we then re-referenced the data to the common average. We did not interpolate any electrode nor correct the EEG signal for the presence of ocular artefacts. The fade-in and the fade-out periods were excluded from the analyses leading to the segmentation of an EEG signal of 60 seconds (corresponding to the display of 600 stimuli). We averaged the signal from all repetitions^3^ of each condition for each participant. We then performed a Fast Fourier Transformation (FFT), and we extracted amplitude spectra for the 128 channels with a frequency resolution (the size of the frequency bins) of 0.016 Hz. Given the topography of our results (see Figure 2a), we merged some electrodes into two regions of interest: we subsequently computed left (A9, A10, A11) and right (B6, B7, B8) occipitoparietal areas of interest.

Based on the frequency spectra, we computed two measures to determine whether and how the brain specifically responded to the deviant frequency during the three manipulated conditions: we first computed the Signal-to-Noise Ratios (SNRs) by dividing each frequency bin by the mean amplitude of their respective twenty surrounding bins (excluding the immediately adjacent bins, and the two most extreme values ^[73, 77, 88:90]^). We used SNRs to depict the frequency spectra and illustrate the topographies of our results (see Figure 2).

We also computed a Z-score to assess the statistical significance of the brain responses to the category change. To do so, for each condition and for each participant, we cropped the FFT spectra around the frequency of interest (1.25 Hz) and its subsequent harmonics up to the seventh (*i.e.*, 1.25, 2.5, 3.75, 5, 6.25, 7.5, and 8.75 Hz) surrounded by their twenty respective neighbouring bins (ten on each side ^[77]^). We summed all cropped spectra and then applied a Z-transformation to the amplitudes. We finally extracted from this Z-transformation the value across the frequency bins of interest. This value represents the brain response specific to the experimental manipulation at 1.25Hz, which can be interpreted as the neural detection of the change of the stimulus category. As a Z-score, a value larger than the threshold of 1.64 (*p* < .05, one-tailed, testing signal level > noise level) indicates a significant response to our experimental manipulation.

## Ethical considerations

All procedures performed in this study were in accordance with the ethical standards of the APA, and with the 1964 Helsinki Declaration and its later amendments or comparable ethical standards. The Ethic Review Panel from the University of Luxembourg approved the methodology and the implementation of the experiment before the start of data collection. Informed consent was obtained from all individual participants included in the study.

## Authors’ contribution

All authors contributed to designing the study; CS and AVR directed the project; MG created the acquisition software; MG and AP collected the data; AP analyzed the data; AP and MG wrote the paper with input from AVR and CS.

## Acknowledgements

The authors would like to thank Aliette Lochy for her advises in data analysis.

## Footnotes

1 For alternative mechanisms underlying the SNARC effect, please also consider the *dual route model* ^[37, 38]^, the *polarity correspondence account* ^[39]^, and the *working memory account* ^[40:45]^.

2 Nevertheless some SNARC studies use tasks not entailing any numerical triggering such as orientation discrimination ^[46]^, plural versus singular nouns ^[47]^, line bisection ^[48]^ or colour discrimination ^[49, 50]^. Significant SNARC effects reported in these studies could therefore be taken as convincing marks of automaticity of numerical magnitude processing.

3 We did not expect that brain responses to deviant odd digits within a sequence of standard even digits would be different than brain responses to deviant even digits within a sequence of standard odd digits in the Parity condition (and similarly for small and large numbers in the Magnitude condition). We thus aggregated all repetitions for these conditions.

